# Substitution Rate Variation in a Robust Procellariiform Seabird Phylogeny is not Solely Explained by Body Mass, Flight Efficiency, Population Size or Life History Traits

**DOI:** 10.1101/2021.07.27.453752

**Authors:** Andrea Estandía, R. Terry Chesser, Helen F. James, Max A. Levy, Joan Ferrer Obiol, Vincent Bretagnolle, Jacob González-Solís, Andreanna J. Welch

## Abstract

Substitution rate variation among branches can lead to inaccurate reconstructions of evolutionary relationships and obscure the true phylogeny of affected clades. Body mass is often assumed to have a major influence on substitution rate, though other factors such as population size, life history traits, and flight demands are also thought to have an influence. Birds of the order Procellariiformes—which encompasses petrels, storm-petrels and albatrosses—show a striking 900-fold difference in body mass between the smallest and largest members, divergent life history traits, and substantial heterogeneity in mitochondrial substitution rates. Here, we used genome-scale nuclear DNA sequence data from 4365 ultraconserved element loci (UCEs) in 51 procellariiform species to examine whether phylogenetic reconstruction using genome-wide datasets is robust to the presence of rate heterogeneity, and to identify predictors of substitution rate variation. Our results provide a backbone phylogeny for procellariiform seabirds and resolve several controversies about the evolutionary history of the order, demonstrating that albatrosses are basal, storm-petrels are paraphyletic and diving petrels nestled within the Procellariidae. We find evidence of rate variation; however, all phylogenetic analyses using both concatenation and multispecies coalescent approaches recovered the same branching topology, including analyses implementing different clock models, and analyses of the most and least clock-like loci. Overall, we find that rate heterogeneity is little impacted by body mass, population size, age at first breeding, and longevity but moderately correlated with hand-wing index, a proxy for wing shape and flight efficiency. Given our results and the context of the broader literature perhaps it is time that we begin to question the prevailing paradigm that one or a few traits largely explain rate variation and accept instead that substitution rate may be the product of weak interactions among many, potentially taxon-specific, variables.

---

Understanding the tempo and mode of molecular evolution is central to evolutionary biology. Molecular substitution rates vary across the Tree of Life but reaching consensus on the factors that influence this phenomenon has proved elusive. Body mass has repeatedly emerged as one of the main drivers of substitution rate (Martin and Palumbi 1993; Mindell and Thacker 1996; Gillooly et al. 2005; Gillooly and Allen 2007) as it is linked to metabolic rate (Brown et al. 2007): smaller animal taxa have higher metabolic rates and increased abundance of free radicals during oxidative metabolism which leads to higher mutation (Brown et al. 2004) and substitution rates (i.e., mutation plus fixation among taxa over time, (Kimura 1968)). Indeed, small-bodied animals often show higher rates of substitution than large-bodied animals (reptiles: Bromham 2002; mammals: Welch et al. 2008; mammals and birds: Nabholz et al. 2013; birds: Jarvis et al. 2014; Weber et al. 2014; Berv and Field 2018). However, other studies spanning a variety of taxa and using both nuclear and mitochondrial DNA have failed to find such relationship (invertebrates: Thomas et al. 2006; birds: Lanfear et al. 2010).

Identifying factors that explain rate heterogeneity is a difficult task because of phylogenetic non-independence of rate estimates (Bromham 2020) and the interaction among multiple life history traits (Simpson 1944; Western and Ssemakula 1982; Brown 1995; Roff 2002). For example, small-bodied taxa tend to have shorter generation times accumulating larger numbers of mutations during many successive rounds of genome replication over a given period of time. Body mass also correlates with population size (Western and Ssemakula 1982), as large taxa tend to have smaller population sizes and experience more genetic drift (Kimura 1968), influencing rates of substitution (Ohta 1972; Lanfear et al. 2014).

Substitution rates can also be influenced by different lifestyles. For example, volant birds spend a large amount of energy in flying (Guigueno et al. 2019). They can increase their flight efficiency by having higher capillary densities in muscles involved in flight (Maillet and Weber 2007), an elongated wing morphology (Dawideit et al. 2009; Garrard et al. 2012) and adopting flight strategies, like dynamic soaring (Bousquet et al. 2017). Despite these adaptations, birds with high flight demands usually show higher substitution rates (Jenni-Eiermann et al. 2014). The hand-wing index (HWI) can be used as a proxy of wing shape and metabolic expenditure (Wright et al. 2014), which is simultaneously linked to flight style.

Rate heterogeneity leads to differences in reconstructed branch lengths that can potentially obscure evolutionary relationships (Felsenstein 1978; Hendy and Penny 1989; Anderson and Swofford 2004; Delsuc et al. 2005). For example, long branch attraction (LBA) is a common artefact arising from extensive rate heterogeneity (Felsenstein 1978), which can be aggravated by sparse taxon sampling (Heath et al. 2008). Although rate of substitution is a fundamental issue in phylogenetics, there is no clear understanding of the impact that rate heterogeneity has in phylogenetic reconstruction (Field et al. 2019), in part due to relatively low variation in body mass, population sizes and life history traits among study species.

Procellariiformes, the most diverse order of oceanic birds, provides an ideal system for examining these factors because of vast differences in body mass (900-fold difference between the largest and smallest; Fig. 1), substantial variation in life history characters and population size. They also demonstrate different flight modes from continuous flapping (e.g. *Pelecanoides* species), wing-propelled diving (e.g. shearwaters), pattering at the surface (e.g. storm-petrels), and dynamic soaring (e.g. albatrosses). In addition to this, there is evidence for mitochondrial molecular rate heterogeneity (Nunn and Stanley 1998). The more than 120 species of Procellariiformes are important marine predators. This group of birds are used as bioindicators of marine productivity and food availability (Nations 2017), but many species are highly endangered – the percentage of threatened procellariiform species is much higher than in Aves overall (>50% in Procellariiformes vs. < 25% in Aves) (Rodríguez et al. 2019). For this reason, correctly understanding their evolutionary relationships has important consequences.

**Fig. 1.**
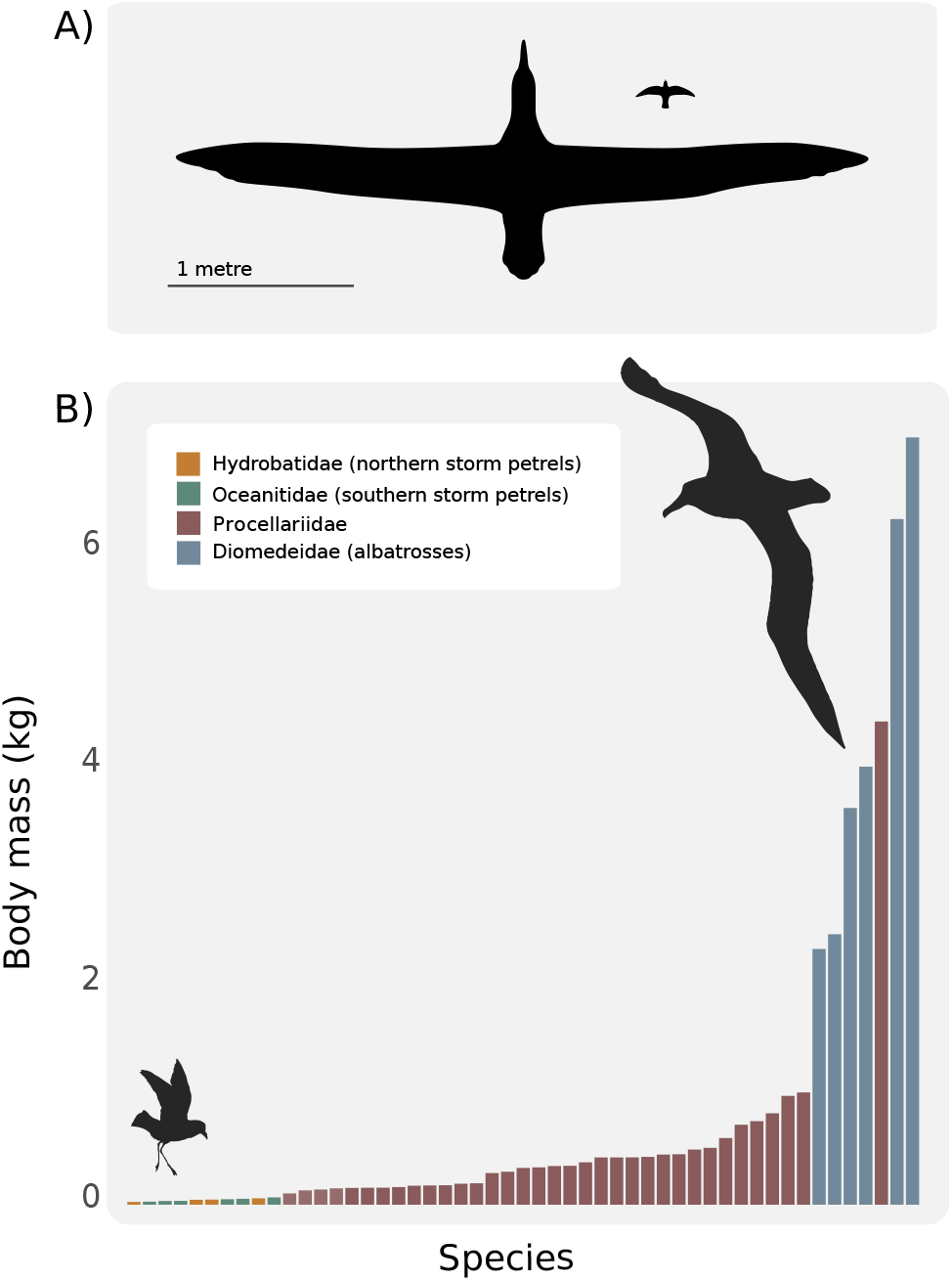
The avian order Procellariiformes includes species with a 900-fold difference in body mass, providing an excellent study system for testing the influence of body mass and life history traits on substitution rate and phylogenetic reconstruction among species. A) Silhouettes of the Least Storm-Petrel (*Oceanodroma microsoma*; wingspan 36 cm) and Wandering Albatross (*Diomedea exulans*; wingspan 3.5 m), to scale. B) Distribution of body mass in various groups of Procellariiformes, which range from 18.2 g to 16.1 kg. The Procellariidae shows the widest variation, largely due to the giant petrels (*Macronectes*), which are unusually large for this family.

Four families are now usually recognised within the Procellariiformes: the albatrosses (Family Diomedeidae), two families of storm-petrels (Families Hydrobatidae and Oceanitidae) and a very diverse group containing gadfly and fulmarine petrels, prions, and diving-petrels (Family Procellariidae) (Warham 1990). The branching pattern of the four families of Procellariiformes remains unresolved (Nunn and Stanley 1998; Kennedy et al. 2002; Hackett et al. 2008; Prum et al. 2015; Reddy et al. 2017). For instance, diving petrels (*Pelecanoides*) were long considered to constitute a separate family (Pelecanoididae) and this classification has received some molecular support (e.g., Sibley et al. 1990; Nunn and Stanley 1998). Conversely, storm-petrels were previously considered to belong to a single family (Hydrobatidae), based on morphological analyses (Livezey and Zusi 2007; Mayr and Smith 2012), but genetic results suggest that they are paraphyletic (Nunn and Stanley 1998; Kennedy et al. 2002; Hackett et al. 2008; Prum et al. 2015). In the past, studies have relied on one or a few mitochondrial markers or taxon-poor datasets (Nunn and Stanley 1998, Kennedy et al. 2002; Penhallurick and Wink 2004; Mayr and Smith 2012). More recently, several higher-level avian phylogenomic studies have included a limited number of procellariform species in their trees (Ericson et al. 2006; Hackett et al. 2008; Prum et al. 2015; Reddy et al. 2017), but because of this sparse taxon sampling little can be concluded about procellariiform relationships. Rate heterogeneity and related differences in branch lengths (as suggested by massive differences in body mass) may be an important factor that has been obscuring the true phylogenetic relationships in this order. In fact, Nunn and Stanley (1998) suggested a correlation between body mass and molecular rate heterogeneity in the mitochondrial gene cytochrome-b in the Procellariiformes. However, differences in life history traits, population sizes and other proxies for metabolic rate were not assessed as possible alternative causes.

Here, we use sequences of 4,365 ultraconserved elements (UCE) from 51 procellariiform species, representing all genera and all major lineages, to reconstruct the deep evolutionary history of this highly diverse order, examine the impact of rate heterogeneity on phylogenetic reconstruction, and analyse the effect of several traits on substitution rate. We find evidence of substitution rate variation across the phylogeny, explicitly exploring the fit of strict and relaxed molecular clocks in a Bayesian framework and also building phylogenies with the most and least clock-like UCE loci. The same topology is consistently recovered, with high support in all nodes, indicating that phylogeny reconstruction with a massive dataset is apparently robust to rate heterogeneity in this avian order. To investigate whether nucleotide substitution rate is correlated with differences in adult body mass, life history traits (age at first breeding [AFB] and longevity), population size, and flight demands (assessed using HWI, as a proxy; Claramunt et al. 2012; Sheard et al. 2020; Duchêne et al. 2021), we implement a Bayesian approach and find that, when controlling for phylogenetic relatedness and other covariates, contrary to expectations, body mass, several life history traits, and population size are only weakly correlated with substitution rate variation, while HWI is moderately correlated. Our results challenge the idea that any single factor can explain a large portion of rate heterogeneity among taxa. We also provide evidence of a robust backbone phylogeny for the Procellariiformes, ending a long-standing debate about their phylogenetic relationships. This phylogeny will allow future work to tackle different aspects of the evolutionary history of Procellariiformes.

## Materials and Methods

### Taxon Sampling

We assembled tissue and blood samples from 51 individuals representing all major lineages within the order Procellariiformes (Table S1). We additionally sequenced three species as outgroups: (i) African penguin (*Spheniscus demersus*), (ii) Maguari stork (*Ciconia maguari*) and (iii) Brown booby (*Sula leucogaster*). Penguins (Sphenisciformes) are the sister order to Procellariiformes, and storks (Ciconiiformes) and boobies (Suliformes) represent more distantly related orders (Prum et al. 2015).

### Library Preparation, Targeted Enrichment of UCEs and Sequencing

We extracted genomic DNA using Qiagen DNeasy Blood & Tissue Kits (Qiagen Inc., Valencia, CA, USA). We quantified all DNA extracts with a Qubit Fluorometer (Life Technologies, Inc.) using the high sensitivity kit and assessed DNA quality by gel electrophoresis. UCE capture and sequencing from our extracts was performed by RAPiD Genomics (Gainesville, FL, USA), as described in Ferrer Obiol et al. (2021). Each pool was enriched using custom-designed probes targeting 5,060 UCEs across tetrapods (Faircloth et al. 2012) and PCR was conducted for 11 cycles to amplify the enriched library. Sequencing was performed on two lanes of an Illumina HiSeq 3000 platform using 100 bp paired-end (PE) sequencing.

### Assembly, Alignment, Trimming and Data Matrices

We evaluated the quality of the raw reads with FASTQC (Babraham Bioinformatics) and then removed Illumina adapter contamination and trimmed low-quality regions with the parallel wrapper script Illumiprocessor (https://github.com/fairclothlab/illumiprocessor). The PHYLUCE package v1.5.0 (Faircloth 2016; https://github.com/faircloth-lab/phyluce) was used for the initial sequence processing stages: We de novo assembled the reads into contigs with Trinity (*assemblo_trinity.py*) (Grabherr et al. 2013) and matched the assembled contigs to UCE probes (*match_contigs_to_loci.py*) (Faircloth et al. 2012). A list containing the UCE loci enriched in each taxon was generated, and then we created a FASTA file with data for each taxon and UCE locus (*get_fastas_from_match_counts.py*).

UCEs were aligned using MAFFT (Katoh et al. 2005; *phyluce_align_seqcap_align.py*) and the alignments were trimmed using a parallel wrapper around Gblocks (*phyluce_align_get_gblocks_trimmed_alignments_from_untrimmed.py*; Castresana 2000). Following alignment, we created three matrices with varying degrees of “completeness”: 95%, 85% and 75% (*align_get_only_loci_with_min_taxa.py*), where “completeness” for a 75% matrix means that all loci were present for at least 75% of the taxa. We calculated the number of informative sites for each alignment (*get_align_summary_data.py* and *get_informative_sites.py*) and evaluated their quality with the TriStats module within the TriFusion package (http://odiogosilva.github.io/TriFusion/; Fischer et al. 2011). We generated PHYLIP-formatted, concatenated alignments based on our final data matrices (*format_nexus_files_for_raxml.py*) for phylogenetic analyses.

For most analyses we used the 75% matrix because it comprises the longest alignment (2,328,288 bp from 4,365 UCE loci) with still a low amount of missing data (10.64%). This matrix contained 294,892 variable sites (12.67%) and 141,901 informative sites (6.09%). The 95% matrix (1,096,476 bp) was exclusively used for the Bayesian partitioned analysis because the length of the other alignments (2,200,911 bp [85% matrix] and 2,328,288 bp [75% matrix]) meant that computational demands were too high.

### Phylogenetic Analyses

#### Partitioning strategies and substitution model testing

We created two partitioning schemes: i) First, we applied a locus-partitioned scheme (partitioning scheme one), which uses each UCE as a separate partition with a potentially separate substitution model and ii) second, we employed an entropy-based partitioning scheme using SWSC-EN (partitioning scheme two; Tagliacollo and Lanfear 2018), which splits each UCE locus into 3 parts – the core and the two flanking regions, and allows different substitution models for each region of each UCE locus. The resulting partitions from each analysis were used as input for PartitionFinder2, which finds the best partitioning schemes by lumping together those partitions that have similar substitution rates and selects the best-fitting substitution model for each subset (Lanfear et al. 2017). We linked the branch lengths and used a relaxed hierarchical clustering algorithm (rcluster method) with the –rcluster-max option set to 5000. We only evaluated the models that RAxML can accommodate (GTR, GTR+G and GTR+I+G) and selected the best scheme using the Akaike information criterion (AIC) with correction for small sample sizes.

### Maximum likelihood analyses

Maximum likelihood (ML) phylogenetic analyses were performed with RAxML v8.0.19 (Stamatakis 2014). We used two approaches: we performed i) a partitioned analysis (using scheme two described above) for the 75% matrix only, and ii) unpartitioned analyses for each of the matrices: 75%, 85% and 95%. Based on PartitionFinder2 results, GTR and GTR+G substitution models were the best fit models for most of the partitions, so for the unpartitioned analyses we ran ML analyses under each of those two models for all three matrices. We performed non-parametric bootstrap replicates using the autoMRE function. Results were visualised using Figtree v1.4.3 (http://tree.bio.ed.ac.uk/software/figtree/; Rambaut 2017).

### Compositional bias and heterotachous evolution

Compositional bias is a type of systematic bias can lead to inconsistent phylogenetic results (Phillips et al. 2004). We calculated the GC content and average base counts for each species and performed a chi-square test to identify potential base compositional bias. We also performed a ML analysis in RAxML using a “RY-coded” matrix that codes bases as either purine or pyrimidine (Woese et al. 1991; Phillips et al. 2004). We used the 75% alignment to generate this matrix and we applied the same settings as above.

To account for heterotachous evolution (rate heterogeneity across sites), we also ran IQ-TREE v.1.6. (Nguyen et al. 2015; https://iqtree.org/) with our 75% matrix and partitioning scheme two. We applied a GTR model with four General Heterogeneous evolution On a Single Topology (GHOST) linked classes (Crotty 2017).

### Bayesian analyses

To estimate phylogenies in a Bayesian framework, we used ExaBayes v1.5 (Aberer et al. 2014). We conducted an unpartitioned analysis on the 75% matrix, with four independent runs each, using default priors and four coupled chains (three heated chains and one cold) for one million iterations and sampling every 500 generations. Convergence was assessed with Tracer v1.7.1 (Rambaut et al. 2018) by examining the estimated sample sizes (ESS>200) and checking the presence of a horizontal trend in traces of the likelihood across generations. The analysis converged and we applied a 10% burn-in. Due to computational demands, the partitioned analysis was performed on the 95% matrix and the locus partitioned scheme (scheme one). Two independent runs for one million generations were carried out and a 10% burn-in was applied.

### Species tree analyses

To conduct species tree analyses, we first used RAxML v.8.0.19, to estimate gene trees for each of the 4,365 UCE loci we recovered, under a GTR+G substitution model and 100 bootstrap replicates. Species trees were calculated without multilocus bootstrapping in ASTRAL-III, as recommended by Sayyari and Mirarab (2016), who showed that internal branch support values are more reliable when bootstrapping is not performed. Due to the high conservation of the UCE core, these markers can have a low number of informative sites. Therefore, we used the AMAS software (Borowiec 2016) to calculate the number of informative sites for each locus, selected the top 10% of loci that had a higher proportion of parsimony informative sites, and then repeated the analysis as described above.

### Lineage-specific rate heterogeneity

To assess the level of rate heterogeneity across the phylogeny, we tested two hypotheses: (i) a null model where all lineages evolved under a constant rate of evolution (i.e. a strict clock) and (ii) a model allowing rates of evolution to be variable among lineages (the alternative hypothesis, an uncorrelated relaxed clock).

First, we conducted two independent sets of analyses in BEAST v1.8.4 (Bouckaert et al. 2014), one with the strict clock model and one with an uncorrelated log-normal relaxed clock model. Due to data set size limitations in BEAST, we created 100 subsets of our data by randomly selecting and concatenating UCE loci sequences so that the total length was between 19,000 and 20,000 bp. For each subset we tested the fit of different substitution models – GTR, GTR+G, GTR+G+I, HKY+G, HKY+G+I and HKY – by calculating Bayes Factors (BFs) and found that HKY best fit our data. We ran an independent BEAST analysis for each subset. Tree topology and clock model (either strict or relaxed) were linked by locus and branch lengths were not, allowing branch length to vary across partitions that way taking into account potential heterotachy. We applied a birth-death process and ran the analyses for 20 million iterations, sampling every 20,000. All replicates converged in fewer than 20 million iterations with all parameters showing an ESS greater than 200. Log files were checked for convergence and adequate ESS with Tracer v1.7.1. We used *Sumtrees* v4.0.0 within the *Dendropy* v4.4.0 python package (Sukumaran and Holder 2010) to summarise all trees into a consensus tree using a Maximum Clade Credibility Topology (i.e. a topology that maximises the product of the clade PPs) and retained clades with a posterior probability (PP) greater than 0.5.

To calculate which clock model provided the best fit for our data, we used a stepping stone/path sampling analysis (Leaché et al. 2014). Marginal likelihood estimates (MLE) were calculated for each subset using 10 million generations and sampling every 10,000 steps. We then calculated each model MLE (strict or relaxed) by combining all the subset output files under each clock. The MLE results allowed us to calculate the BFs and compare the best-fitting model. We calculated ln(BF) using the BF=2 x (MLE(model 1) – MLE(model 2)) and evaluated the strength of the support using the Kass and Raftery (1995) framework. We also assessed for rate heterogeneity by extracting the Coefficient of Variance (CoV) from the BEAST runs. The CoV shows how much the sequence data deviate from clock-likeness, and values below 0.1 indicate strong evidence for a strict clock (Drummond and Bouckaert 2015).

Second, to further test the robustness of our topology to rate heterogeneity, we used SortaDate (Smith et al. 2018) to create two subsets (most clock-like and least clock-like) each of which approximately contained 25% of our total number of loci and using the same piece of software we calculated the root-to-tip variance of each subset, spanning around 1 million bp. We ran RAxML analyses on the two subsets using the settings previously described.

Third, we ran BEAST again, this time using fossil calibrations to examine substitution rates among linages. We used three datasets: 1) randomly selected UCE loci, 2) the most clock-like loci and 3) the least clock-like loci. Each subset comprised approximately 30 loci spanning 20,000 bp. We fixed the topology using the Exabayes tree as a starting tree and set an uncorrelated lognormal clock. Using an off-set-exponential distribution, we specified minimum bounds for seven calibrated nodes, including the root of the tree. The node age for the most recent common ancestor (MRCA) of Procellariiformes and Sphenisciformes is based on *Waimanu manneringi* (60.5 Ma), a giant extinct penguin found in New Zealand by Slack et al. (2006). We also calibrated the following crown nodes: 1) Crown Diomedeidae, 2) Crown Oceanitidae, 3) Crown *Pelecanoides*, 4) Crown *Puffinus*, 5) Crown *Calonectris*, and 6) Crown *Ardenna*; the last three fossils were selected following Ferrer Obiol et al. (2021) (see Supplementary Information for further information).

### Phylogenetic comparative analyses

We collected body mass, AFB, and population size data from Brooke (2004), The Handbook of the Birds of the World (del Hoyo et al. 2011) and the primary literature (see Tables S3-S5), longevity from Myhrvold et al. (2015) and HWI from Sheard et al. (2020). When female and male body mass were available we used the mean of both measurements. We also considered analysing clutch size, but there is no variation within the Procellariiformes for this trait.

To explore how body mass and other life history variables were correlated with each other while accounting for phylogenetic non-independence, we used the R package *phylopath* v1.0.0 (van der Bijl 2018). We tested nine models that included several combinations of possible relationships among body mass, AFB, longevity, population size, and HWI. We performed 1000 bootstrap replicates and selected the best model by using the *best* function in phylopath.

We performed analyses with a birth-death speciation prior and an autocorrelated relaxed clock in Coevol 1.4b to explore how the variables included in the phylogenetic path analyses were correlated with substitution rate (Lartillot and Poujol 2011; Lartillot and Delsuc 2012). As Coevol 1.4b runs are computationally intensive, we selected and concatenated approximately 150,000 bp from UCEs that contained a high percentage of parsimony informative sites according to AMAS. We used the ExaBa-yes topology for the starting tree and used the fossil calibrations described in the previous section. We ran the MCMC chain for approximately 1000 iterations until convergence was reached (ESS > 200). We ran four replicates for each analysis and discard 10% of the iterations as burn-in. Because of missing data for certain variables, like population size or longevity, we decided to also run a separate analysis with only AFB and body mass.

GC-biased gene conversion is known to affect avian genomes (Nabholz et al. 2011; Weber et al. 2014) especially those with large population sizes and small body mass (Weber et al. 2014). We explored whether this could be the case in our group of birds by creating two subsets, each yielding a concatenated alignment of approximately 150,000 bp: 1) we selected UCEs based on high GC-content (top 10%) and 2) those with lowest GC-content (bottom 10%). We re-ran Coevol 1.4b with the same parameters, number of independent analyses, iterations and burn-in as specified above.

## Results

### UCE Sequence Data

The mean number of reads per sample was 1.98 million (Standard deviation (SD) = 80,536) and we recovered a total of 4,926 UCE loci. Individual UCE alignments had a mean length of 576.73 bp (Table 1; SD = 10.5). The frequency of missing data was very low in all matrices, with values lower than 14% (Table 1). GC content was homogeneous across taxa (Table S2, χ^2^=0.001, p-value=1.0).

**Table 1.** Summary of the 95%, 85% and 75% data matrices.

|  | 95% Matrix | 85% Matrix | 75% Matrix |
| --- | --- | --- | --- |
| **Min. taxa per locus** | 51 | 45 | 40 |
| **Alignment length (bp)** | 1096476 | 2200911 | 2328288 |
| **Missing data (%)** | 5.88 | 10.64 | 13.34 |
| **GC content (%)** | 38.17 | 39.18 | 39.15 |
| **Loci (n)** | 1908 | 4048 | 4365 |
| **Min. len. of UCE loci (bp)** | 245 | 203 | 203 |
| **Max. len. of UCE loci (bp)** | 1205 | 1522 | 1522 |
| **Mean len. of UCE loci (bp)** | 574 | 544 | 534 |

### Phylogenetic Analyses

For partitioning scheme one, in which data were initially demarcated by locus, PartitionFinder suggested there were 2709 (75% matrix) and 1696 distinct partitions (95% matrix). The best-fitting substitution model for partitions was GTR+I+G, followed by GTR (Table S6). The entropy-based partitioning analysis (SWSC-EN, partitioning scheme two) identified 2,927 distinct data partitions in the 75% matrix, all of which followed the GTR+G model.

The topologies recovered from different tree building methods were essentially identical. All the RAx-ML results – 75%, 85%, 95% unpartitioned analyses implemented with a GTR+G substitution model, as well as the 75% partitioned (partitioning scheme two), and the analysis with the RY-coded matrix – yielded the same fully resolved topology with all nodes supported with bootstrap values of 100 (Fig. 2). IQ-TREE yielded the same topology as well, though two nodes (the *Ardenna-Calonectris* node and the *Thalassoica-Daption* node) received bootstrap support of 99 and 98, respectively, rather than 100 (Fig. S2). Trees from the BI analyses – 75% unpartitioned and 95% partitioned (partitioning scheme one) – show the same topology as the ML trees with all nodes showing PPs of 1.

**Fig. 2.**
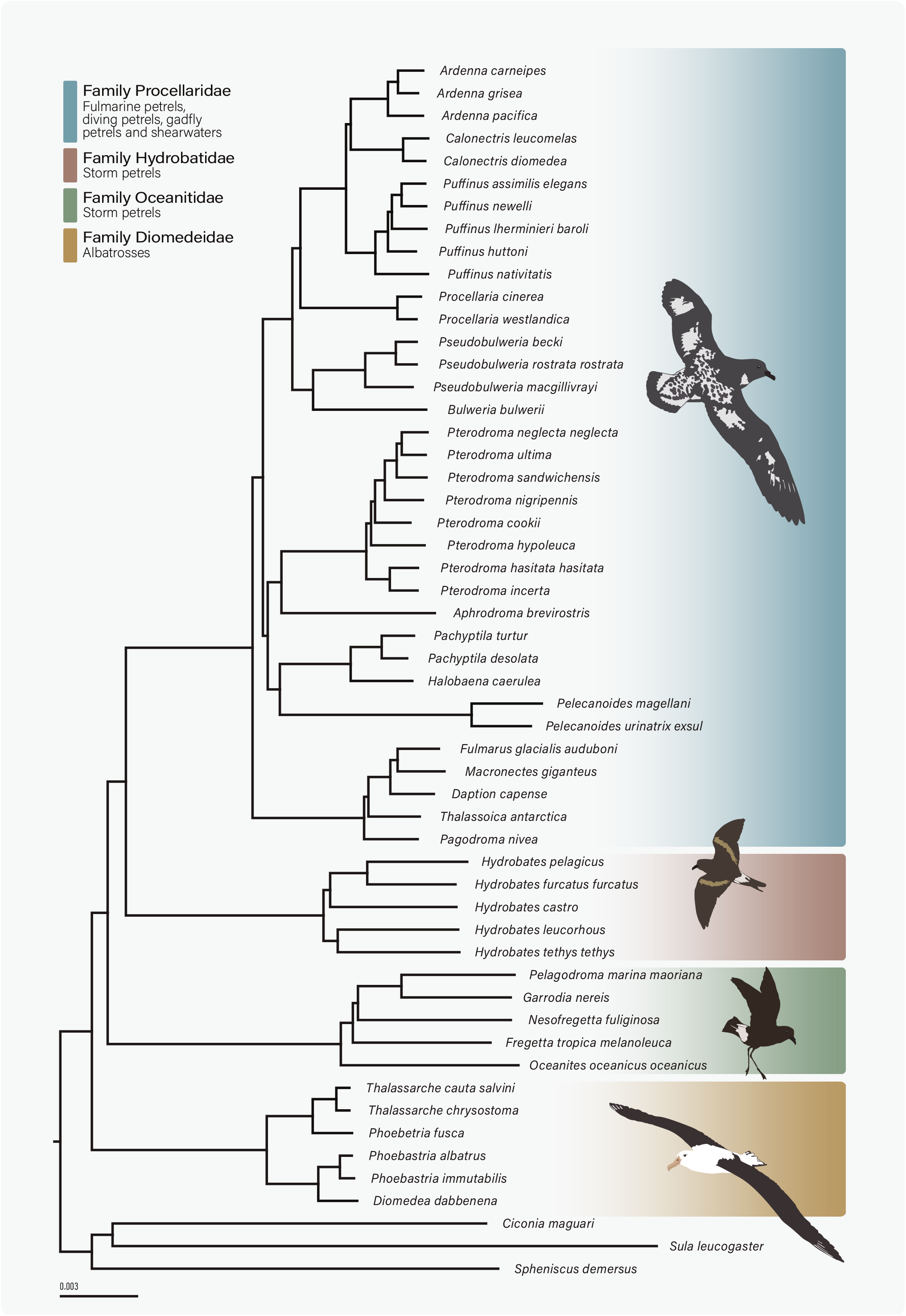
Best-fit phylogeny for the Procellariiformes from the ExaBayes analysis. All posterior probabilities are 1.0. All ML and Bayesian analyses, using different substitution models and different partitioning schemes, produced the same topology.

In all analyses, the albatrosses (Diomedeidae) are sister to all other ingroup taxa (Fig. 2). The two families of storm-petrels (Oceanitidae and Hydrobatidae) did not form a monophyletic group and constitute instead a paraphyletic group including Procellariidae, which appears as the sister group of Hydrobatidae. Within Procellariidae, the fulmarine petrels (*Fulmarus*, *Macronectes*, *Daption*, *Thalassoica*, and *Pagodroma*) are sister to the rest of the group, which forms two clades. In one clade, the diving-petrels (sometimes considered a separate family), prions (*Pachyptila* spp.) and the monotypic *Halobaena caerulea* form a monophyletic group sister to the gadfly petrels (*Pterodroma* spp.). In the other clade, *Bulweria* and *Pseudobulweria* are sister genera and form a monophyletic group sister to the shearwaters and the petrels of the genus *Procellaria*, which are sister taxa.

We used two approaches for building species trees: we built gene trees for (i) every UCE locus in our most dataset and (ii) the top 10% most informative UCE loci. Species trees applying both methods show the same topology as in the ML and BI analyses (Fig. S3). The support values correspond to local PPs as described in (Sayyari and Mirarab 2016). As with other analyses, support was very high over most of the tree, although two nodes (the node uniting the three genera of shearwaters and the node uniting the diving petrels and the gadfly petrels) received lower support (all UCEs: 0.77 and 0.86; top 10: 0.67 and 0.91, respectively).

### Lineage-specific Rate Heterogeneity

To investigate the impact of among-lineage rate variation, we first applied strict and uncorrelated relaxed clock models in BEAST for 100 randomly selected approximately 20,000 bp subsets of our data. We found strong support (BF=151; ln(2*BF) = 5.71) for the uncorrelated lognormal relaxed clock model (MLE = −42933) over the strict clock model (MLE = −43084), suggesting that different lineages in our topology experience different nucleotide substitution rates. The CoV, extracted from the BEAST runs, shows how much the sequence data deviate from clock-likeness, and values below 0.1 indicate strong evidence for a strict clock (Drummond and Bouckaert 2015). We find a mean CoV of 0.5 (95% Highest

Posterior Density (HPD)=0.3-0.7) indicating strong deviations from clock-likeness. The consensus tree resulting from both clock models showed the same topology obtained by all other approaches (ML, ExaBayes and ASTRAL) (Fig. S4). Further, for the second test, the topologies recovered using both the top 25% clock-like UCE loci and the 25% least clock-like loci were the same as well, although both subsets had a slightly lower support at some nodes. Interestingly, the clock-like subset shows lower support for the node uniting the three groups of shearwaters (55) and the node uniting the diving petrels and the gadfly petrels (75), whereas the non-clock-like subset shows a support of 60 and a 100, respectively.

For the third test, we performed independent fossil-calibrated BEAST runs on random, clock-like and non-clock-like subset. All three time-calibrated trees strongly rejected clock-like evolution (random UCEs: mean CoV=1.24, 95% HPD=0.83-1.66, clock-like subset: mean CoV=1, 95% HPD=0.60-1.78, non-clock-like subset: mean CoV =1.15, 95% HPD=0.70-1.74). When we plotted the substitution rate onto the tree, we found that most variation was at internal nodes, especially those leading to the different subgroups within the Procellaridae, and those leading to the shearwaters (Fig. 3).

**Fig. 3.**
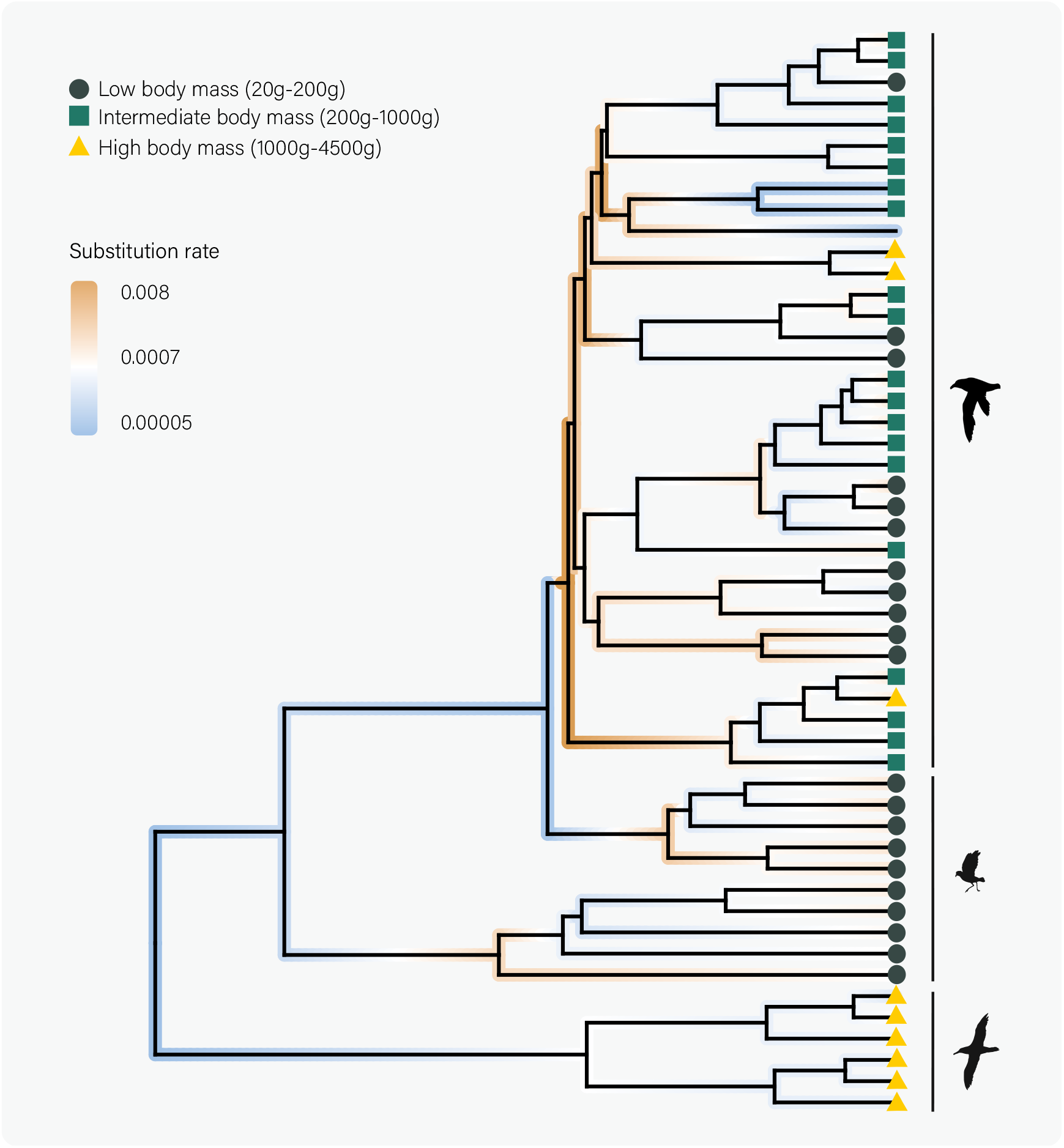
BEAST tree showing substitution rates mapped onto the phylogenetic tree. The highest substitution rates are mainly present at internal nodes and at terminal branches of small-bodied birds.

### Correlates of substitution rate variation

The Procellariidae showed the widest variability in body mass with some species weighing as little as 100 g (*Bulweria Bulwerii*) and others as much as 4.5 kg (*Macronectes giganteus*) (Fig. 1; Tables S3-S4). For AFB and population size, the storm-petrels had the largest populations and breed relatively early in life while the albatrosses have smaller populations and breed later. Longevity and HWI were also variable across families but showed lower variation overall (Table S5). The best model extracted from the phylogenetic path analysis included relationships between AFB and HWI (R^2^=−0.43), body mass (R^2^=0.55), population size (R^2^=−0.38), and longevity (R^2^=0.15). The best model also included relationships between HWI and body mass (R^2^=−0.31) and longevity (R^2^=0.13). However, only for the relationship between AFB and body mass the confidence intervals did not overlap with zero (Fig. S6). Coevol 1.4b performs pairwise tests integrating partial correlations among other variables while controlling for phylogenetic relationships. AFB and body mass were weakly negatively correlated with substitution rate after controlling for all the other variables (R^2^= −0.01, PP=0.77, and R^2^= −0.02, PP=0.7, respectively. Table S7). HWI showed a moderate negative effect on substitution rate, with a high posterior probability (R^2^=−0.18, PP=0.95). Longevity showed a small effect on substitution rate (R^2^=−0.05, PP=0.74) and population size showed the opposite trend but the effect has the same intensity as longevity (R^2^=0.04, PP=0.74). Note that posterior probabilities for all traits, except for HWI, are relatively low. The analysis that only included AFB and body mass, and had more data points, recovered a slightly higher correlation coefficients (AFB R^2^= −0.15, PP =0.99; Body mass R^2^= −0.07, PP=0.83, Table S8). When analysing UCE loci selected based on high and low GC-content we found the same trend as in the model including all variables with random UCEs (Tables S9-S10; Fig. S7).

## Discussion

We hypothesised that given the extreme differences in body mass and variation in life history traits in the Procellariiformes, extensive rate heterogeneity would be present and could potentially bias phylogenetic inference. We did uncover relatively high rate heterogeneity, at the higher end of the range of CoV found in other orders of birds (i.e. Passeriformes) (Nguyen and Ho 2016). Despite the presence of rate heterogeneity, all tree-building methods, including those implementing variable and strict clock models as well as the most clock-like and least-clock like loci, produced the same robust topology. We tested for correlations among substitution rate and several variables, yet we only found weak correlation coefficients. Our results indicate that body mass, population size, HWI, longevity and generation time effects do not explain a large proportion of the rate variation in this order of birds, which challenges the commonly held assumption that only a small set of traits is largely responsible for driving substitution rates.

### Correlations with Substitution Rate

Several hypotheses have been proposed to explain substitution rate heterogeneity among lineages and particular hypotheses have received support in different taxonomic groups (Martin and Palumbi 1993; Mooers and Harvey 1994; Galtier et al. 2009; Garcia-Porta et al. 2019; Duchêne et al. 2021). Smaller bodied organisms tend to have higher metabolic (Nagy 2005) and substitution rates (Gillooly et al. 2005), including some species of Procellariiformes (Brown and Adams 1984). Other early work suggested that body mass but not AFB was correlated with the mtDNA substitution rate in procellariiform seabirds (Nunn and Stanley 1998), but rigorous data analysis frameworks that account for correlations between traits, phylogenetic relatedness, and explicit incorporation of fossil calibrations were not available at that time.

Our Coevol 1.4b results revealed very weak correlation coefficients between substitution rate and body mass. Previous studies have found correlations of different strengths between body mass and substitution rate. For example, using the same method for their regression models and a similar number of data points, Weber et al. (2014) reported that approximately 10% of the variation of substitution rate in the class Aves could be explained by body mass alone and 15% in the case of Berv and Field (2018). Our results suggest a much smaller percentage of the variation explained by body mass alone (2%; Table S7). This lower value may be influenced by the fact that our study uses a lower taxonomic rank, though Procellariiformes has the most variation in body mass of any bird order. The presence of missing data could also influence our results. In fact, when we ran an analysis with only body mass and AFB, to reduce the amount of missing data, we find that body mass still only accounted for 7% of the substitution rate variability (Table S8). Since body mass is inversely correlated with metabolic rate, these results seem to suggest that the importance of metabolic rate in driving substitution rate in this order might not be as important as previously thought. In line with our results, other studies have sometimes also reported very weak correlations between substitution rate and body mass (Mooers and Harvey 1994; Lanfear et al. 2010; Lourenço et al. 2013).

The time-calibrated tree (Fig. 3) reveals that the highest substitution rates are present at internal branches, especially those branches that lead to the different Procellariidae groups. The Coevol 1.4b ancestral reconstruction of body mass at the internal nodes of the Procellariidae reveals intermediate-sized birds with very high substitution rates, suggesting that body mass might not be driving substitution rates. Rates may differ at different points in time because species do not keep the same body mass or life history traits throughout their evolution (Bromham 2011). This has often been overlooked as many studies have performed phylogenetic independent contrasts incorporating traits only at the tips and not including ancestral state reconstructions. That said, Figure 4 suggests that the negative correlation is strongest when taking into account both the internal branches and the tips, though subtler when considering the tips alone. Furthermore, ancestral state reconstructions can sometimes have large confidence intervals, which can also influence precision of point estimates (Pagel et al. 2004).

**Fig. 4.**
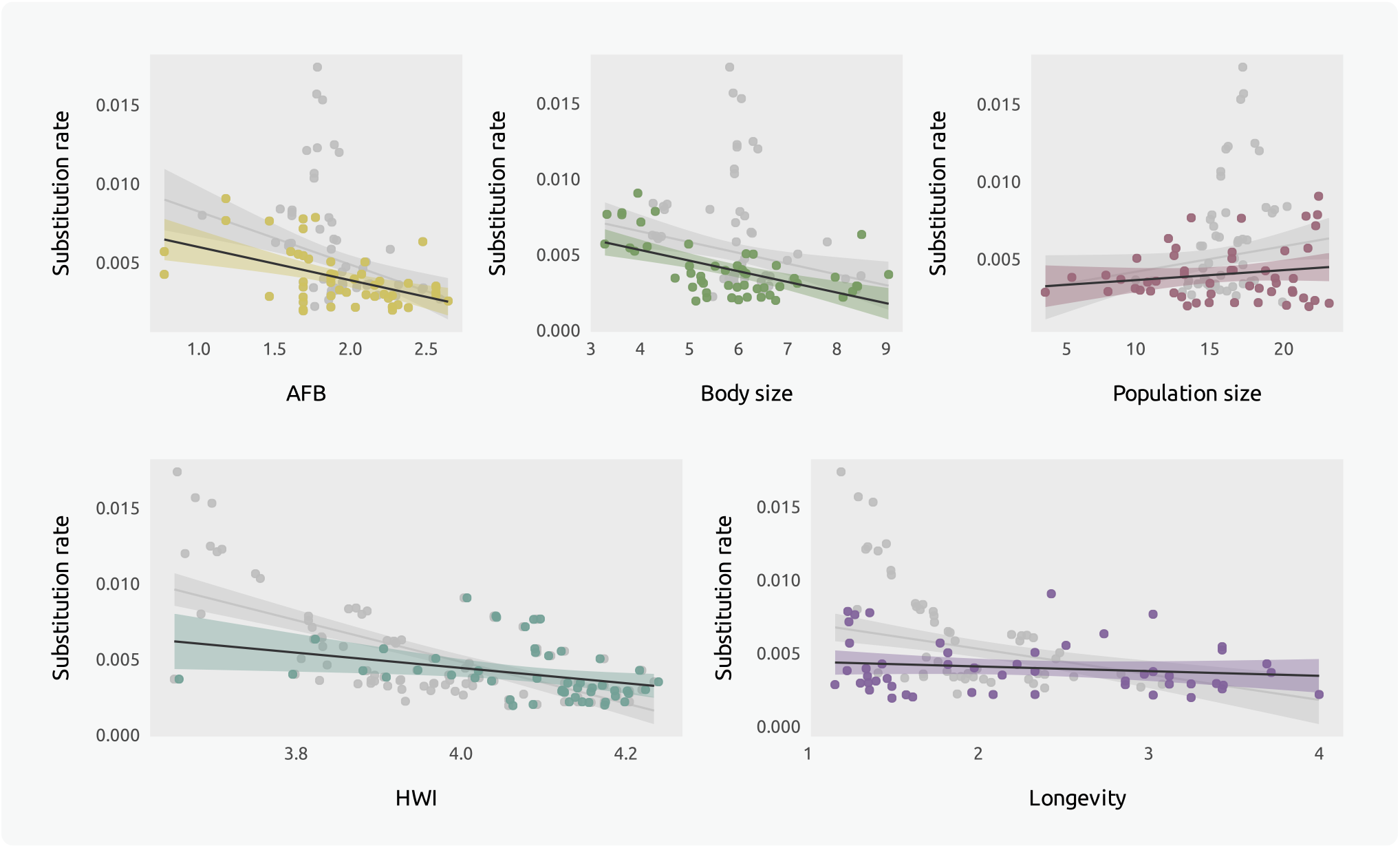
Substitution rates (calculated as the estimate of the instant value of the rate at the node or tip by Coevol 1.4b) plotted against each of the variables included in the model (log-transformed). The highlighted points represent the trait at the tips and the grey points represent reconstructed traits at internal nodes. The black line and coloured 95% credibility intervals (CI) represent the fit line for the tips while grey fit lines and 95% CI represent the mean trend (tips+internal nodes). All variables are log-transformed. Trends tend to be similar at the tips and internal nodes, but stronger at internal nodes. AFB: Age at first breeding; HWI: Hand-wing Index.

Like body mass, HWI can also be considered to be a proxy for metabolic rate (Ward et al. 2001). It has been used as a proxy for flight efficiency (Duchêne et al. 2021), where those species that are highly dispersive species and efficient flyers show higher HWI. We find a moderate negative correlation between HWI and substitution rate that explains 18% of the variation. These results suggest that those birds with low HWI values might fly less efficiently, expend more energy, have a higher metabolism, and produce a higher amount of nuclear substitutions. Contrary to our results on body mass, this suggests that metabolic rate may indeed be important. Body mass has usually been correlated with basal metabolic rate in other species, but Procellariiformes spend relatively little time resting on land —they spend most of their lives at sea, are capable of traveling exceptionally long distances without rest (Shaffer et al. 2006), and may even sleep while flying, although this idea is debated (Rattenborg 2017). The metabolic rate during flight might be more important than basal metabolic rate in this order of birds. Energetic expenditure during flight is not only determined by morphology, but also by physiological and behavioural adaptations to flight, which we are not necessarily reflected fully by HWI, therefore future research could investigate the role of flight in greater detail.

Generation time, longevity and population size have also previously been proposed to have an important effect on rates of nuclear substitution (Chao and Carr 1993; Welch et al. 2008; Thomas et al. 2010). AFB, a proxy for generation time, has the same effect as body mass on substitution rate, only explaining 1% of the variation in the analyses that included all variables, but 15% in the analysis that only included AFB and body mass. In the latter model, we had a denser dataset, which reduced the uncertainty when reconstructing ancestral traits. Furthermore, it is likely that we obtained a higher correlation coefficient because we were not taking into account the correlations between AFB and all the other covariates that were included in the complete model. Nunn and Stanley (1998) found that AFB did not explain substitution rate variation in Procellariiformes, but they also pointed out that they had very few data points for performing independent contrasts. Estandia (2019) used phylogenetic generalised least squares to explore the relationship between body mass, AFB and substitution rates in Procellariiformes, reporting much higher correlation coefficients (AFB: R^2^=0.64 and body mass: R^2^=0.56), but ancestral state reconstructions were not included and a different estimate of substitution rate was used.

Longevity was hypothesised to be an important factor driving substitution rate variation in birds by Berv and Field (2018), but they suggested that it was actually not a good predictor after accounting for correlations among other covariates. However, in fishes longevity alone can explain up to 16% of the variation in substitution rates (Hua et al. 2015). While obtaining reliable longevity data for seabirds is challenging, especially for those species that live on remote islands, longevity is known for being highly variable in procellariiform seabirds (Wasser and Sherman 2010). Some albatross species show an average lifespan of 40 years or more while diving petrels can live as little as 6 years. We find that 5% of the variability in substitution rates can be explained by longevity in this group, though this may be influenced by missing data.

Population size is often thought to be positively correlated with substitution rate because of body mass - large-bodied organisms tend to have small populations, long generation time, lower metabolic rate, and hence lower substitution rates (Ohta 1972; Western and Ssemakula 1982; Lanfear et al. 2014). However, the phylogenetic path analysis for this group of birds reveals a very weak correlation between population size and body mass and given that the relationship between population size and substitution rate is slightly stronger than with body mass, it is then unlikely that the small effect of population size is due to body mass effects. We note that census size can be a weak proxy for effective population size in some cases (Nabholz et al. 2013) and that severe, recent population bottlenecks have occurred in procellariform species due to human impacts such as habitat loss and introduced predators (Croxall et al. 2012). It is likely that the true relationship between population size and substitution rate has been and will remain obscure.

GC-biased gene conversion can influence the relationship between population size and substitution rate (Bolívar et al. 2019). GC-rich regions have higher mutation and recombination rates (Kiktev et al. 2018), which can influence substitution rates, especially in lineages with large populations and short generation times (Weber et al. 2014). We find that the overall trend indicates comparable relationships between the GC-rich dataset and the GC-poor dataset (Tables S9-S10). These results suggest that GC-biased conversion is likely not an essential factor driving substitution rate heterogeneity in this group. There is, however, an alternative explanation that proposes that as population size increases, the prevalence of natural selection can increase faster than the production of new mutations, which can produce higher advantageous substitution rates, independent of body mass and GC-biased gene conversion (Lanfear et al. 2014).

For all the traits studied here, our results suggest that the relationship with substitution is stronger when considering reconstructed ancestral traits at the nodes in addition to those at the tips (Fig. 4). A possible explanation for this pattern would be that elevated substitution rates during speciation might have a more important impact on substitution rates than body mass or any of the other tested traits, as recently proposed by Janzen et al. (2020). Indeed, in our phylogeny internal branches with high substitution rates are associated with periods of rapid speciation (Ferrer Obiol et al. 2021). However, we remain cautious when interpreting these results, due to the high uncertainty that comes with ancestral reconstructions (Pagel et al. 2004), especially when phylogenetic sampling is sparse.

The fact that many studies reveal varying strengths in the relationships between life history traits, genome characteristics and substitution rates suggests a complex interaction among factors, which likely differs across taxonomic groups (Gillman and Wright 2013). In our study, no single factor explains a large proportion of variation in substitution rates. Therefore, perhaps it is time that we begin to question the prevailing paradigm and propose that substitution rate may be the product of the interaction among many variables, potentially including environmental variables (Rolland et al. 2016) and speciation rates (Janzen et al. 2020).

### Genome-wide Data Yield Robust Topologies for the Procellariiformes Despite the Presence of Rate Heterogeneity

Systematic or non-random errors (e.g. LBA, compositional and rate heterogeneity) are well-known problems in phylogenetics (Felsenstein 1978). A number of solutions have been proposed to overcome the potential bias of among-rate-branch-variation (Carruthers et al. 2020). However, because rate heterogeneity is not as extensive at low taxonomic levels (e.g., genus level), where a large number of phylogenetic studies are concentrated, its consequences during phylogenetic tree construction have not been studied in depth. Given the striking differences in body mass (Fig. 1) as well as variation in life history traits, we hypothesised that rate heterogeneity would be present among lineages and potential LBA could be the reason of the lack of resolution and low support observed in previous studies of the Procellariiformes phylogeny.

Our ML and Bayesian trees support the hypothesis that there is rate heterogeneity across the procellariiform phylogeny. When comparing strict and relaxed clock models, BFs strongly support substitution rate variation across lineages, rejecting the clock-like evolution hypothesis. Our CoV values from this analysis, which measure rate variation, are among the highest values found in the literature for an order of birds (Nguyen and Ho 2016; Berv and Field 2018). Further, the fossil-calibrated BEAST analysis strongly rejected clock-like evolution, with the largest rate heterogeneity occurring between internal and terminal branches. Studies on a range of other taxa across different taxonomic scales have also demonstrated the presence of rate heterogeneity (e.g. Pereira and Baker 2006; Patané et al. 2009; Eo and DeWoody 2010; Beaulieu et al. 2015), suggesting that it may be a widespread issue and may affect phylogenetic reconstruction on a broad scale.

Despite the presence of extensive rate heterogeneity, we recover a highly consistent and well-supported topology using multiple analytical approaches. Although our results are congruent with those from other genome-scale studies, we provide much greater resolution through sequencing 51 species (vs. a maximum of 8 species in other genomic studies) and including representatives of all genera, major lineages, and at least five species from each family. Our results strongly support a single topology (i) with the albatrosses (Diomedeidae) as the sister group to all other taxa, (ii) a paraphyletic relationship for the two families of storm-petrels (Hydrobatidae and Oceanitidae), (iii) Hydrobatidae as sister taxon to the Procellariidae, and (iv) the diving-petrels (*Pelecanoides*) placed within Procellariidae, rather than as a distinct family.

This topology also clarifies long-debated relationships of some procellariiform species and genera. For example, the Kerguelen petrel (*Aphrodroma brevirostris*) has been described as a “taxonomic oddball”, as evidenced by its taxonomic history: it was first placed within the gadfly petrels (*Pterodroma* spp.), then placed in a monotypic genus (*Lugensa* or *Aphrodroma*), and later considered to be closely related to the shearwaters (Nunn and Stanley 1998; Kennedy and Page 2002). This species is sister to the gadfly petrels in our trees. Another example is the genus *Procellaria*: some studies have suggested that *Procellaria* and *Bulweria* form a monophyletic group sister to the shearwaters (Nunn and Stanley 1998). However, we find Procellaria to be sister to the shearwaters and that *Bulweria* and *Pseudobulweria* are sister to the clade consisting of the shearwaters and *Procellaria*, thus supporting the findings of Kennedy and Page (2002).

LBA is one consequence of rate heterogeneity and a frequent cause of inaccurate phylogenetic inferences (Anderson and Swofford 2004; Bergsten 2005; Brinkmann et al. 2005; Philippe et al. 2005). The family Hydrobatidae (northern storm-petrels) demonstrates a long branch, and this family has repeatedly been reconstructed as sister to the rest of the order in previous studies (Nunn and Stanley 1998; Kennedy and Page 2002; Hackett et al. 2008). Those results may have been influenced by LBA, which is even more likely given the relatively sparse datasets (both in terms of taxon sampling and in the length and number of loci) employed. Despite the large body mass difference between the storm-petrels and the albatrosses we found that body mass has a weak association with substitution rate and is unlikely to cause LBA in this phylogeny, because the short-branched albatrosses arise as the basal group.

The presence of short internal branches indicative of periods of rapid speciation has also been found to obscure phylogenetic reconstruction (e.g. Alda et al. 2018; Ferrer Obiol et al. 2021). Rapid speciation can lead to incomplete lineage sorting (ILS), which may result in the loss of phylogenetic signal and conflicting topologies (Rokas and Carroll 2006; Whitfield and Lockhart 2007; Wiens et al. 2008; Philippe et al. 2011; McCormack et al. 2013; Suh et al. 2015). Concatenation-based methods can be less accurate than coalescent-based approaches in the presence of ILS (i.e. Bayzid and Warnow 2013; Edwards et al. 2016). In this study, we used the summary method ASTRAL-III which is statistically consistent under the multi-species coalescent. Our ASTRAL species trees show the same topology as recovered in the ML and Bayesian analyses, although with somewhat lower support in some nodes. One node with differing support in the species tree analysis is the node uniting the three species of Ardenna and the two species of *Calonectris* shearwaters, with support of 0.77 in the species tree compared to full support in the ML and Bayesian trees (Fig. S3 and Fig. 2). This problematic short internal branch also emerged in Ferrer Obiol et al. (2021), where it was demonstrated that discordance in the shearwater topology was mainly driven by high levels of ILS due to rapid speciation. The branches leading to this node also show a high substitution rate, which agrees with the proposal that nucleotide substitution rates are high during periods of multiple and rapid divergence events (Janzen et al. 2020). The time-calibrated tree places the divergence of each of the three shearwater clades (*Ardenna*, *Calonectris* and *Puffinus*) during the Middle Miocene, between 11.5 Ma and 12.5 Ma, relatively soon after the Middle Miocene Disruption, when a series of extinctions were occurring due to a severe drop in the global temperatures and re-establishment of the East Antarctic Ice sheet (Pearson and Palmer 2000). After extinction periods, niches often become available allowing species to radiate (Simpson 1944; Raup 1994; but see Hoyal Cuthill et al. 2020), which might explain the shearwater rapid radiation and ILS.

Another node with slightly lower support in the species tree than in concatenation-based methods is the node uniting *Pterodroma* + *Aphrodroma* with the clade comprising the prions and the diving petrels (*Pelecanoides*), which has full support in the ML and Bayesian analyses but a PP=0.86 in the species tree. As discussed above, lower support might be the result of rapid speciation events, which can generate patterns of phylogenetic incongruence, sometimes due to ILS at short internal branches (Pamilo and Nei 1988; Rosenberg and Nordborg 2002). In some previous phylogenies (Nunn and Stanley 1998; Kennedy and Page 2002) it was hypothesised that the diving petrels were a separate family, Pelecanoididae, because it was sister to other Procellariidae. It is likely that the small amount of mitochondrial DNA sequence previously used, together with the long branch leading to *Pelecanoides*, contributed to placing *Pelecanoides* as sister to rather than within the Procellariidae as recovered in our analyses.

## Conclusion

Our work suggests that large datasets may overcome issues related to among-lineage-rate-variation; however, this should be tested in other taxonomic groups using other types of data. Disparities in body mass, life history traits, and several other factors have been proposed to drive rate heterogeneity, but we find no support for the hypothesis that any of these variables is largely responsible for substitution rate variation alone. Rather, we find that many factors explain a small amount of rate variation, including potentially high substitution rates during rapid speciation events.

## Data accessibility

All raw sequence data are archived on GenBank under the accession number PRJNA749710. Code is available here: https://github.com/andreaestandia/procellariiform_backbone.

## Supplementary material

Supplemental text on fossil calibrations, supplementary figures S1-S7, supplementary tables S1-S10, alignments and phylogenetic trees are available from the Dryad Digital Repository: http://dx.doi.org/10.5061/dryad.

## Acknowledgements

We thank the following people and institutions for providing samples for this study: Gary Nunn; Jeremy Austin; Sharon Birks, Burke Museum of the University of Washington; Chris Milensky, National Museum of Natural History, Smithsonian Institution; Paul Sweet, American Museum of Natural History; Donna Dittmann, Museum of Natural Science of the Louisiana State University; Mark Robbins, Biodiversity Institute of Kansas University; and Hadoram Shirihai. We also thank Daniel Field and Mike Bottery for discussions on data analysis, and Nilo Merino Recalde for his help with the figures and feedback.

