## Supplementary text and figures for "Substitution Rate Variation in a Robust Procellariiform Seabird Phylogeny is not Solely Explained by Body Mass, Flight Efficiency, Population Size or Life History Traits"

**This pdf file includes:**

Supplementary Information Text

Figures S1-S7

**SUPPLEMENTARY INFORMATION TEXT**

### Fossil calibrations

The fossil record of Procellariiformes is sparse when compared with other bird orders, especially its sister order Sphenisciformes (Ksepka & Clarke 2010, Olson 1985c). There are, however, some fossil Procellariiformes that are both robustly dated and identified and therefore suitable for fossil calibrations. Our justification of these fossils, below, follows best practices described by Parham et al. (2012) where possible. For all calibration points only a minimum age was set with no upper constraint specified, except for the root of the tree.

#### 1. Node between Sphenisciformes/Procellariiformes

**Minimum age:** 60.5 Ma
**Maximum age:** 61.5 Ma
**Taxon and specimen:** *Waimanu manneringi* (Slack et al. 2006); CM zfa35 (Canterbury Museum, Christchurch, New Zealand), holotype comprising thoracic vertebrae, caudal vertebrae, pelvis, femur, tibiotarsus, and tarsometatarsus.
**Locality:** Basal Waipara Greensand, Waipara River, New Zealand.
**Phylogenetic justification:** *Waimanu* has been resolved as the basal penguin taxon using morphological data (Slack et al. 2006), as well as combined morphological and molecular datasets (Ksepka et al. 2006, Clarke et al. 2007). Morphological and molecular phylogenies agree on the monophyly of Sphenisciformes and Procellariiformes (Livezey & Zusi 2007, Prum et al. 2015). *Waimanu manneringi* was previously used by Prum et al. (2015) to calibrate Sphenisiciformes, and see Ksepka & Clarke (2015) for a review of the utility of this fossil as a robust calibration point.
**Age justification:** The top of the Waipara Greensand marks the Paleocene-Eocene boundary, and calcareous nanofossils further constrain age of this fossil to 60.5-61.6 Ma (Cooper 2004, Slack et al. 2006, Ksepka & Clarke 2015). The younger species *Waimanu tuatahi* dated to ~58-60 Ma is represented by multiple specimens that overall comprise almost a complete skeleton (Slack et al. 2006), providing further evidence of the presence of Sphenisciformes in the Paleocene after this time.

#### 2. Diomedeidae (basal node)

**Minimum age: 30 Ma**
**Taxon and specimen:** *Tydea septentrionalis* Mayr and Smith 2012a; IRSNB Av 94 (Institut royal des Sciences naturelles de Belgique, Belgium), holotype comprising partial humeri, partial ulna, partial carpometacarpi, partial phalanx proximalis digiti majoris, extremitas cranialis and partial scapula, partial coracoids, partial scapus claviculae.
**Locality:** Boom Formation, Belgium.
**Phylogenetic justification:** The position of *T. septentrionalis* in Diomedeidae is supported by shared derived morphology of the humerus, carpometacarpus, and coracoid, as well as large size relative to other procellariiform families, and is strongly differentiated from the extinct Pelagornithidae which were other seabirds that lived at this time and were similar in size (Mayr & Smith 2012a). Despite strong similarities to extant albatrosses, several plesiomorphic features define *T. septentrionalis* as a stem lineage outside the crown group (Mayr & Smith 2012a).
Several specimens pre-dating *T. septentrionalis* have been assigned to Diomedeidae. From the Middle Eocene, *Murunkus subitus* was assigned to Diomedeidae based on only a single carpometacarpus (Panteleyev & Nessov 1987), and further material would be preferable to firmly establish its affinities (Mayr & Smith 2012a). An incomplete tarsometatarsus from the late Eocene was also assigned to Diomedeidae (Tambussi & Tonni 1988), but this placement has been questioned (Mayr & Smith 2012a). Acosta Hospitaleche & Gelfo

(2017) described the genus *Notoleptos* from the Late Eocene of Antarctica, with tentative placement in Diomedeidae. An incomplete furcula from the late Oligocene of New Zealand, described as *Manu antiquus* and compared to Diomedeidae in its description (Marples 1946), has been considered to have been misidentified (Mayr 2009). Considering the questions remaining over the placement of these potentially older specimens, and that *T. septentrionalis* predates all other described fossil Diomedeidae taxa, such as the late Oligocene *Diomedavus knapptonensis* (Mayr & Goedert 2017) and early Miocene genus *Plotornis* (Mayr & Pavia 2014), this specimen is considered here the most robust calibration point for Diomedeidae at present.
**Age justification:** Fossils were found in the Boom Formation, sediments of which were deposited in a shallow open sea in the southwestern part of the North Sea Basin (Vandenberghe et al. 2004, Mayr & Smith 2012a). The deposits of the Boom Formation are dated to 30-31 Ma (Abels et al. 2007).

**3. Oceanitidae (basal node)**

**Minimum age:** 5 Ma
**Taxon and specimen:** *Oceanodroma (Hydrobates) hubbsi* Miller 1951; No. 39979 (University of California Museum of Paleontology, California, USA), holotype comprising skull, vertebrae, pelvis, and left leg.
**Locality:** South of Capistrano Beach, Capistrano Formation, California, USA.
**Phylogenetic justification:** The position of this specimen in *Hydrobates* (the whole Oceanodroma genus has now been transferred to Hydrobates, del Hoyo et al 2014) is well-supported by measurements of the skull, bill, femur, tibia, and tarsus, which place it close to, or in between, *H. melania* and *H. homochroa*, both extant taxa breeding on the coast of California (Miller 1951). *H. hubbsi* is distinguished from other members of Hydrobates by larger bones of the foot and relative length of the femur (Miller 1951). Other than *H. hubbsi*, Howard (1978) described a partial tarsometatarsus as an indeterminate species of *Hydrobates* from the late Miocene at Laguna Niguel in California, although further material would be preferable to confirm this placement.
**Age justification:** Miller (1951) described the Capistrano Formation as middle Miocene in age, however, more recent works indicate that the site near where this fossil was collected encompasses the boundary of the Miocene and Pliocene (see Deméré and Berta 2005 for review), and as such 5 Ma is used as a suitable minimum age. Although the assignment of a specimen predating *H. hubbsi* to *Hydrobates* by Howard (1978) is based on limited material, it supports the presence of the genus earlier in the Miocene.

**4. *Pelecanoides* (basal node)**

**Minimum age:** 16 Ma
**Taxon and specimen:** *Pelecanoides miokuaka* Worthy et al. 2007; MNZ S.42.431 (Museum of New Zealand Te Papa Tongarewa, Wellington, New Zealand), holotype comprising a partial humerus.
**Locality:** Lower Bannockburn Formation, Manuherikia Group, Otago, New Zealand.
**Phylogenetic justification:** This fossil specimen is distinct from other Procellariidae genera due to several derived features shared with extant members of the diving-petrel genus *Pelecanoides*, and is similar in size to modern *P. georgicus* (Worthy et al. 2007). Worthy et al. (2017) report the discovery of a further humerus referable to this taxon. This is the oldest fossil specimen referred to *Pelecanoides*, predating a specimen described from the middle Miocene (~11-13 Ma; Scofield et al. 2006) and specimens from the Pliocene (Olson 1985b).
**Age justification:** This specimen is from the lower Bannockburn Formation, Manuherikia Group, a lacustrine deposit from Otago in New Zealand, of which the stratigraphy and paleogeography has been detailed (Douglas 1986). The age of the formation, dated to 16-19 Ma, is known from palynological evidence (Mildenhall & Pocknall 1989, Pole & Douglas 1998).

**5. *Puffinus* (basal node)**

**Minimum age:** 15.2 Ma
**Taxon and specimen:** *Puffinus mitchelli* Miller 1961; No. 58184, (University of California Museum of Paleontology, Berkeley, California, USA), holotype comprising a partial humerus. *Puffinus priscus* Miller 1961; No. 58185 (University of California Museum of Paleontology, Berkeley, California, USA), holotype comprising a partial humerus.
**Locality:** Sharktooth Hill, Temblor Formation, California, USA
**Phylogenetic justification:** These two species were described by Miller (1961) and are strongly placed in *Puffinus* due to the distinctive flattened humerus. *P. inceptor* was also described from the same locality (Wetmore 1930), although Miller (1961) considered it to differ from all extant and fossil *Puffinus* sufficiently to warrant a separate subgenus at least, and further material would be preferable to confirm or dispute its taxonomic placement.
Some specimens predating these two species have been described and assigned to *Puffinus*, but are controversial. The early Oligocene *Puffinus raemdonckii* was originally assigned to Laridae (van Beneden 1871), and later transferred to *Puffinus* (Brodkorb 1962). This specimen is most likely procellariiform, but the holotype has been lost and without re-examination or further material cannot reliably be classified (Olson 1985c, Mayr & Smith 2012b). The identification of early Miocene *Puffinus arvernensis* (Milne-Edwards in Shufeldt 1896) has been questioned (Olson 1985c). *Puffinus micraulax* was described from the Hawthorne Formation, Florida, USA and dated to the early Miocene (Brodkorb 1963). However, the Hawthorne Formation consists of several separate formations ranging in age from the early Miocene to early Pliocene. Formations of the early Miocene are rarely exposed near to where *P. micraulax* was found (Scott 1988), and it is likely to date to the middle Miocene at earliest and possibly as recently as the Pliocene. In contrast to these other taxa, the combination of strong taxonomic identification and dating (see below) make *P. mitchelli* and *P. priscus* the best candidates for calibrating *Puffinus*.
**Age justification**: The bonebeds at Sharktooth Hill, where these specimens were found, are dated to 15.2-15.9 Ma using magnetostratigraphy and biostratigraphy (Pyenson et al. 2009).

**6. *Calonectris* (basal node)**

**Minimum age:** 13.8 Ma
**Taxon and specimen:** *Calonectris kurodai* Olson 2009; USNM 237220 (Smithsonian Institution National Museum of Natural History, Washington, D.C., USA) holotype comprising a partial humerus.
**Locality:** Calvert Formation, Westmoreland County, Virginia, USA.
**Phylogenetic justification:** This fossil specimen, and associated paratypes, share several features with extant *Calonectris* species which distinguish it from other genera within Procellariidae and from the other shearwater genera, *Puffinus* and *Ardenna* (Olson 2009). The humerus is considerably smaller than modern *Calonectris* species, contrasting with all other, younger, fossil taxa of the genus from the Pliocene (Olson 1985a, Olson & Rasmussen 2001) and Pleistocene (Olson 2008), possibly suggesting a stem position of *C. kurodai*.
**Age justification:** The holotype of *C. kurodai* was found in Calvert Formation Bed 14, which is dated to the Langhian Age of the middle Miocene (Ward & Andrews 2008, Olson 2009). As such, the end of the Langhian Age is used as a minimum age.

**7. *Ardenna* (basal node)**

**Minimum age:** 13.8 Ma
**Taxon and specimen:** *Puffinus (Ardenna) conradi* Marsh 1870; ANSP 13360 (Academy of Natural Sciences of Drexel University, Philadelphia, Pennsylvania, USA), holotype comprising a partial humerus.
**Locality:** Calvert Formation, Calvert County, Maryland, USA.
**Phylogenetic justification:** This specimen was originally described as *Puffinus conradi* (Marsh 1870), but some members of Puffinus were transferred to Ardenna by del Hoyo et al (2014). Wetmore (1926) examined

the holotype and described it as differing only in minor details from modern *Ardenna gravis* and of the same size and proportions. Kuroda (1954), using skeletal characters of extant and extinct shearwater species, also supported the placement of this species in *Ardenna*. This species predates other known fossil examples of *Ardenna* (Chandler 1990, Tennyson & Mannering 2018) and is used to calibrate the genus.
**Age justification:** The Calvert Formation of Maryland is dated to the middle Miocene, with the earliest deposits dating to the Langhian Age (Ward & Andrews 2008). As such, the end of the Langhian Age is used as a minimum age.

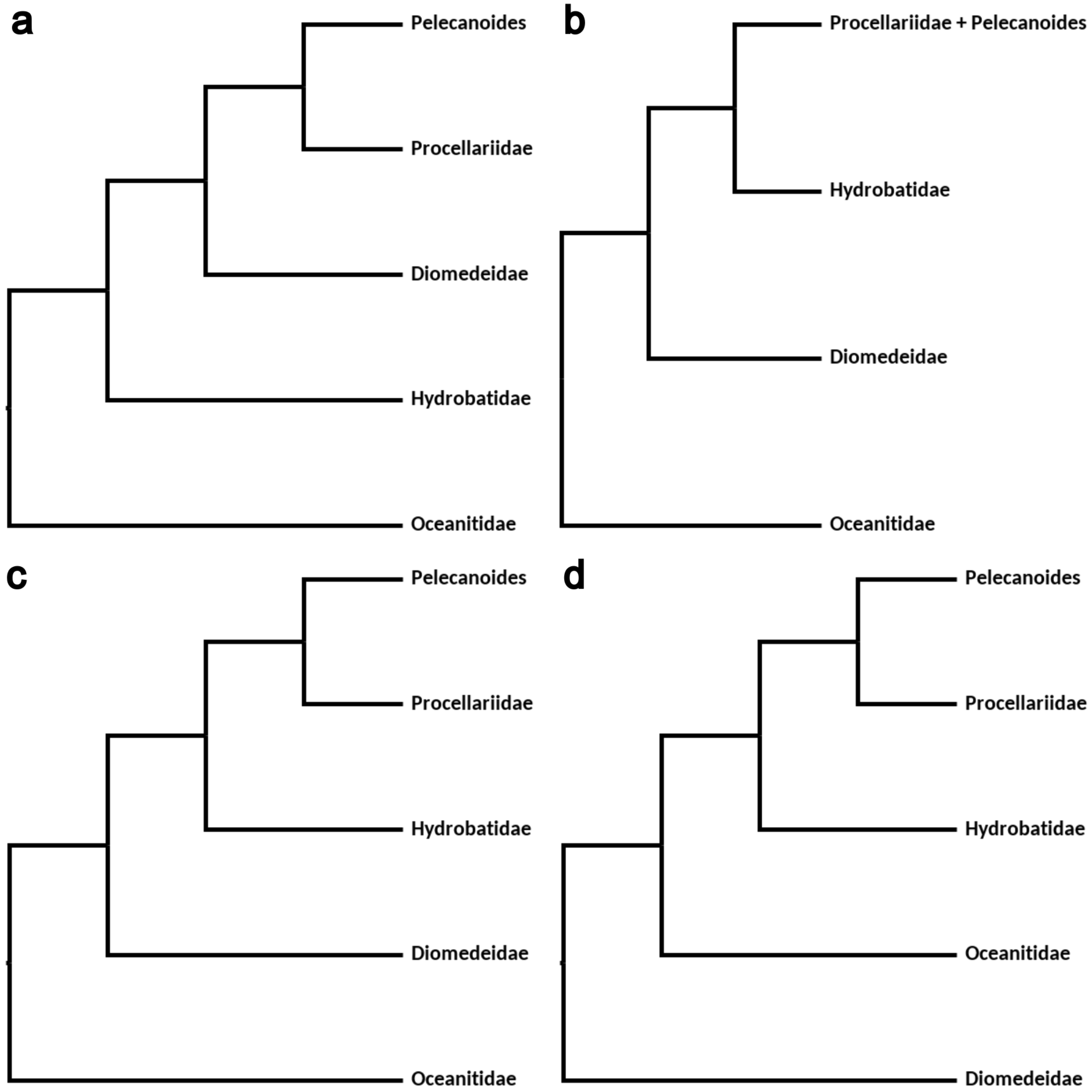


**Figure S1.** Most relevant previously published phylogenetic relationships among the main procellariiform lineages according to (a) Nunn and Stanley 1998, (b) Kennedy and Page 2002 (c) Hackett et al. 2008 and (d) Prum et al. 2015 and Reddy et al. 2017.

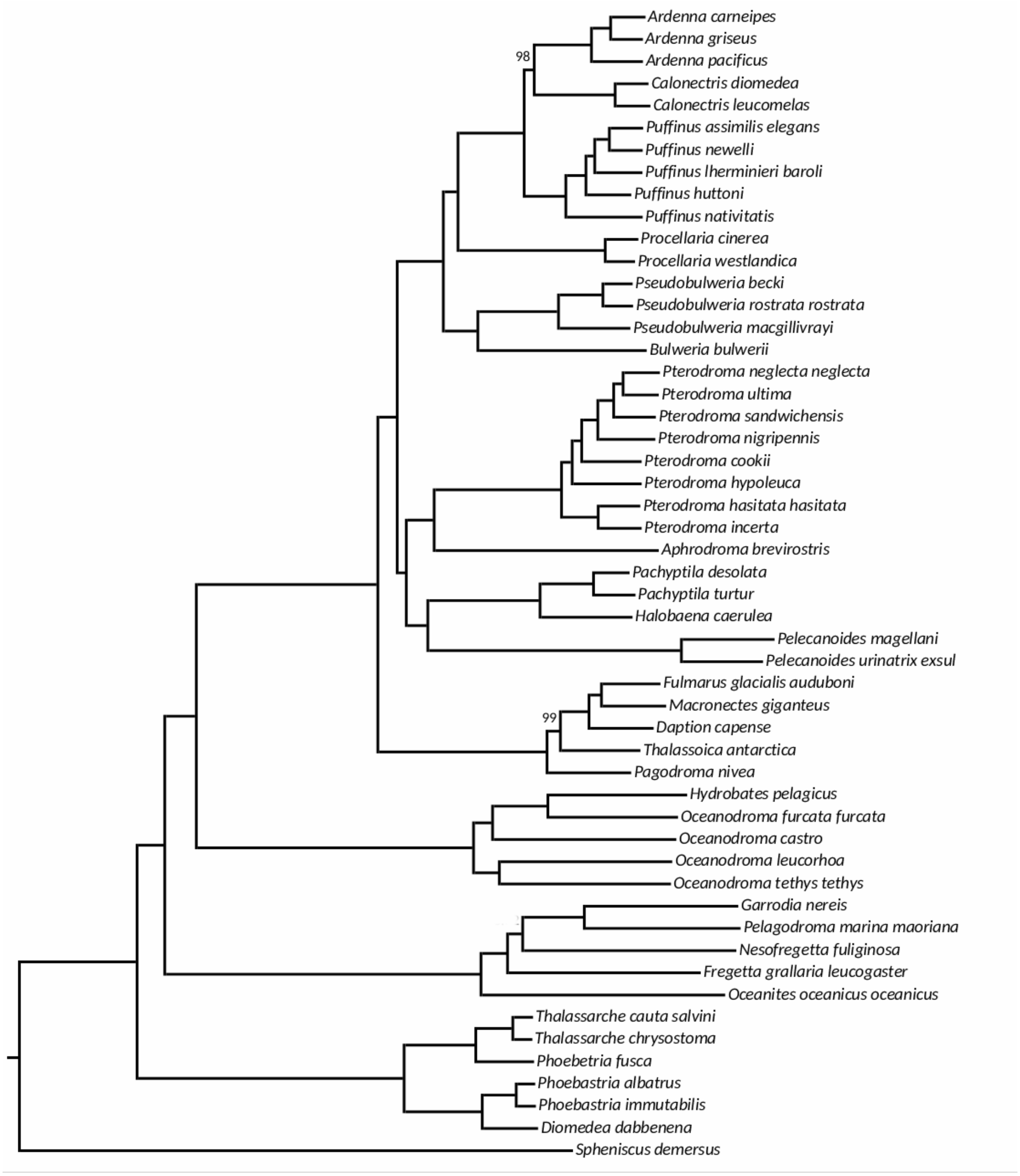


**Figure S2.** Topology resulting from IQ-tree. All nodes are supported by values of 100, except where indicated.

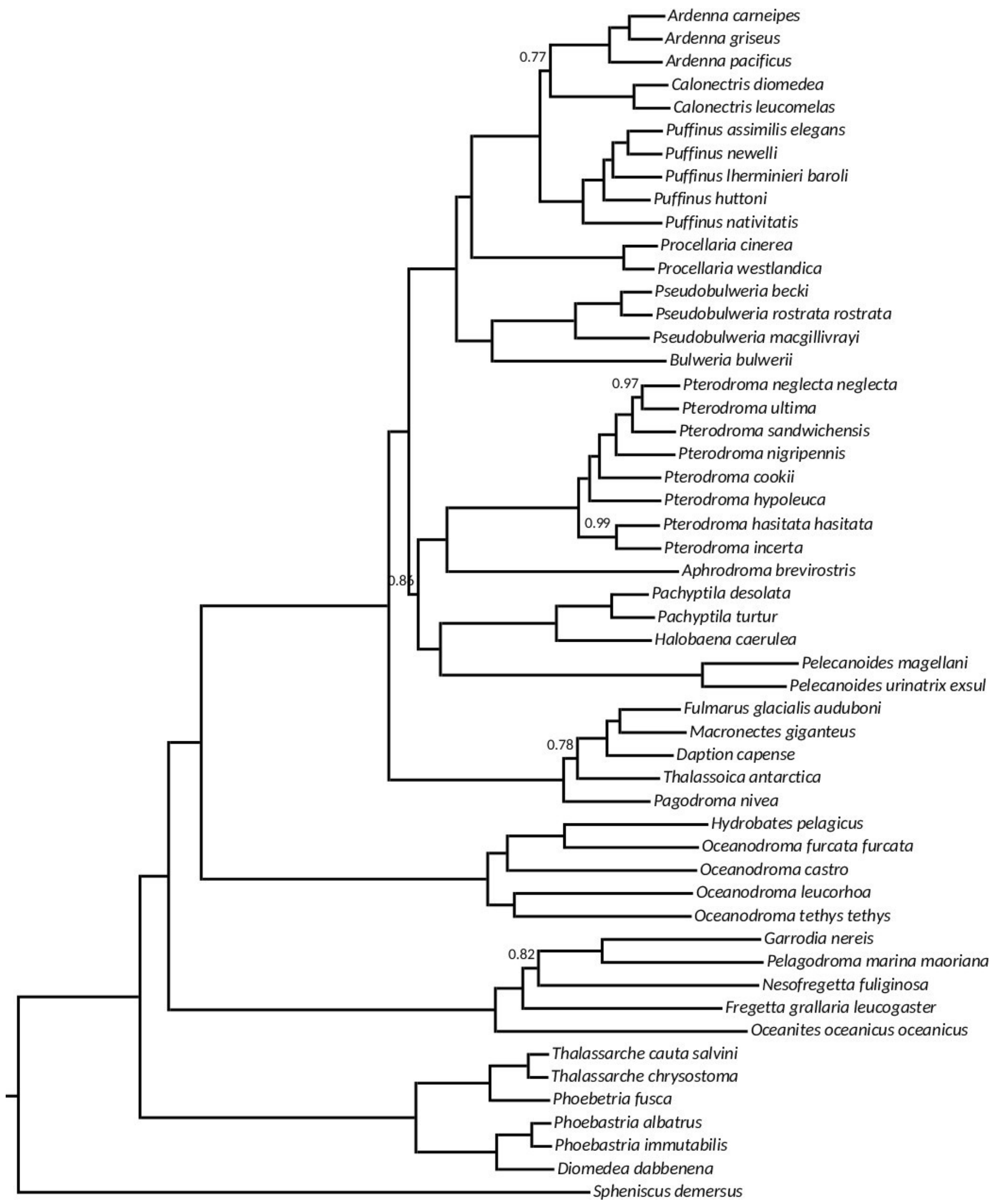


**Figure S3.** Species tree (ASTRAL-III). All nodes supported by values of 1 except for those indicated in the figure.

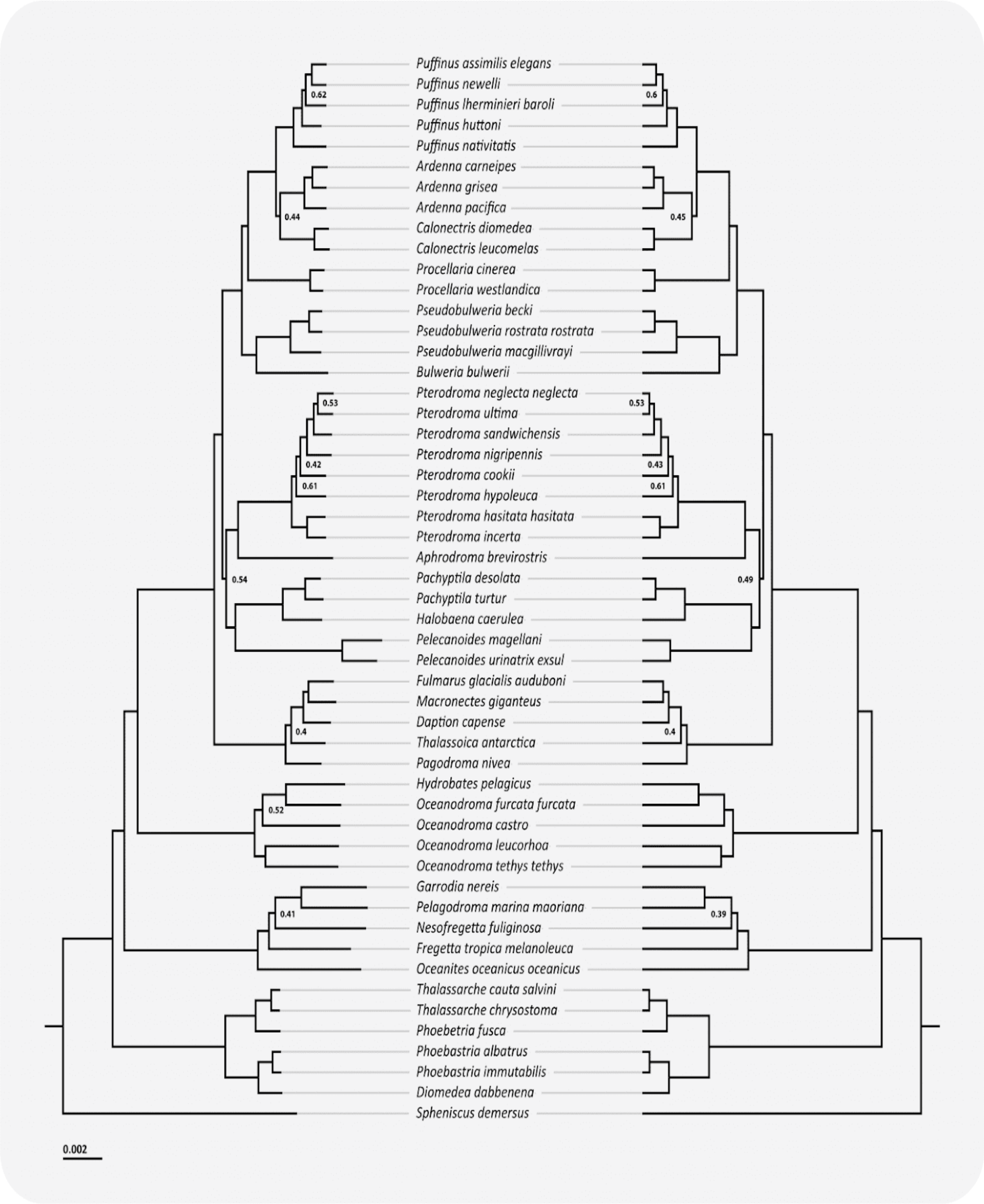


**Figure S4.** Comparison between the BEAST phylogeny under a relaxed clock model (left) and a strict clock model (right). The topology remains the same, but note the differences in branch lengths, particularly the short branches within the Diomedeidae (Thalassarche, Phoebetria, Phoebastria, and Diomedea) and the long branch leading to *Pelecanoides*. All nodes are supported by at least a posterior probability of 0.7 unless otherwise indicated.

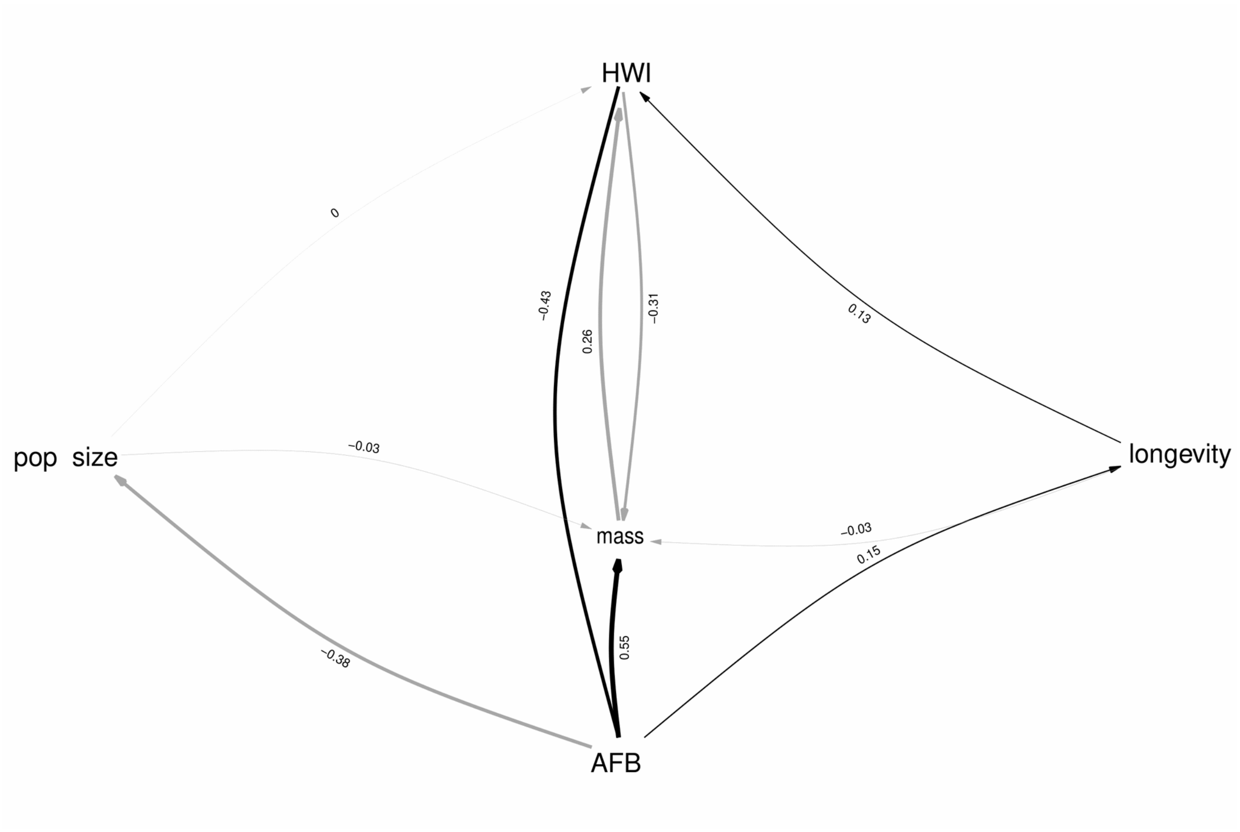


**Figure S5.** Best-fitting phylogenetic path model.

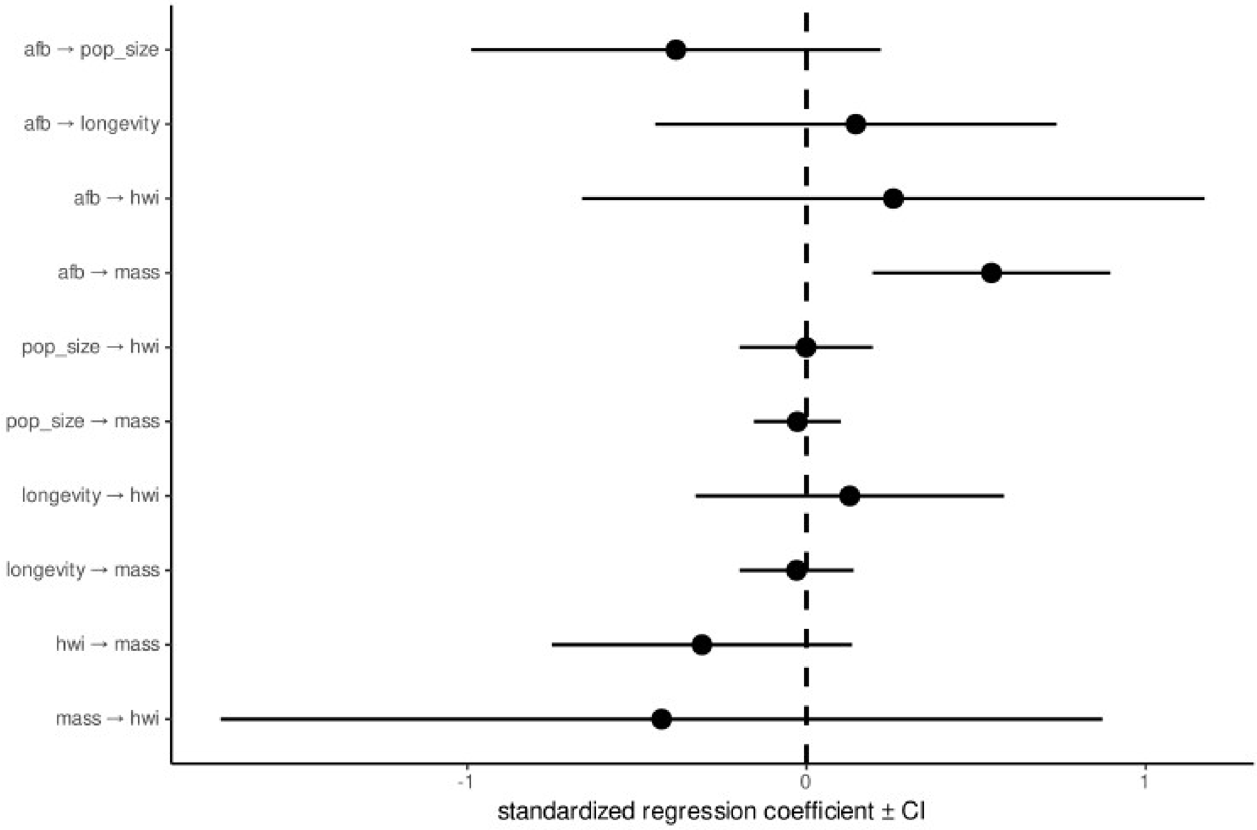


**Figure S6.** Phylogenetic path analysis results. The only significant correlation is between AFB and body mass, as the confidence interval does not overlap zero.

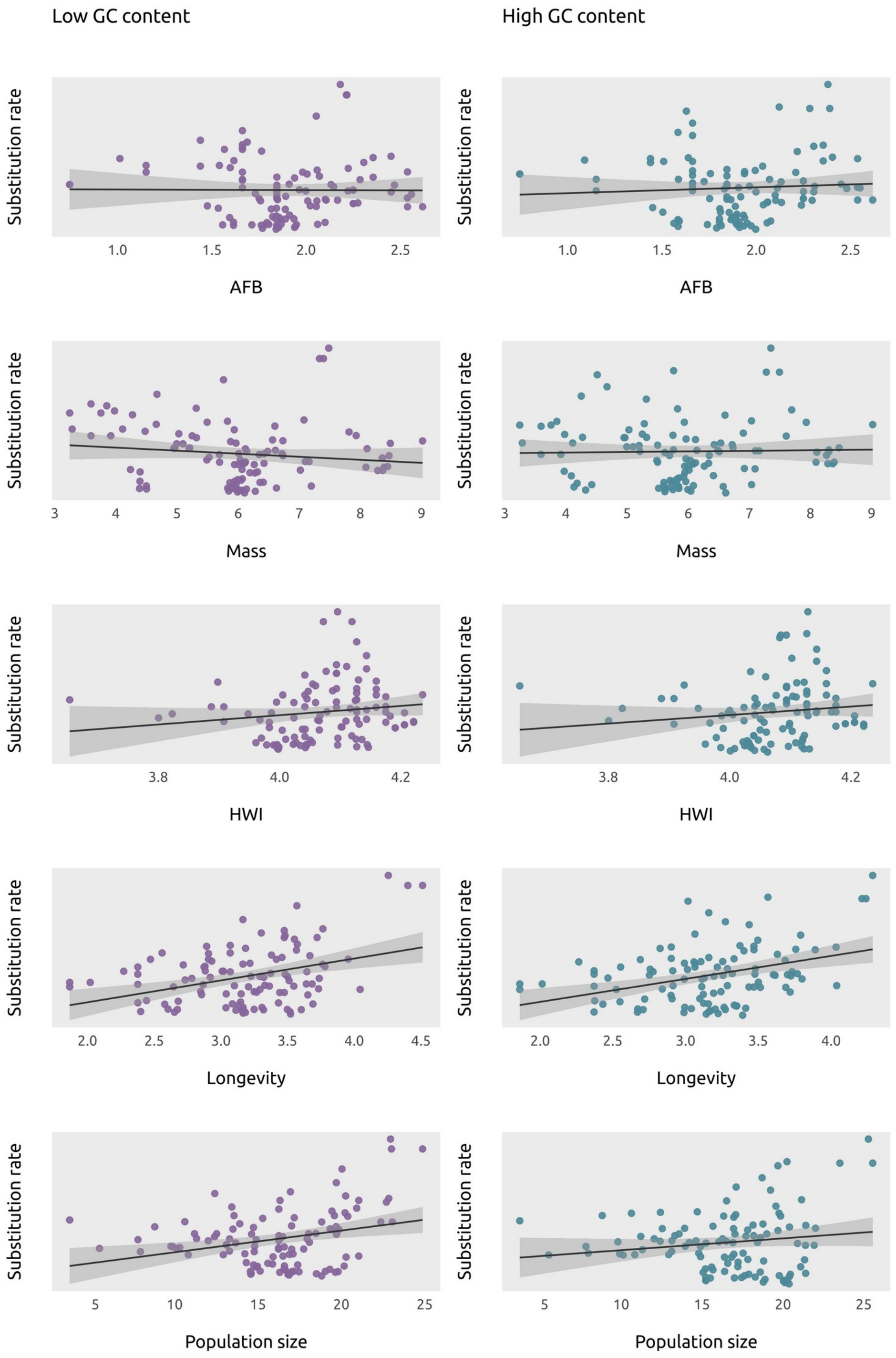


**Figure S7.** Substitution rates (calculated as the estimate of the instant value of the rate at the node or tip by Coevol 1.4b) plotted against each of the variables included in the model (log-transformed) with either only high-GC content or only low-GC content. We find similar strengths in the relationships between both datasets.
